# SIMLR: a tool for large-scale single-cell analysis by multi-kernel learning

**DOI:** 10.1101/118901

**Authors:** Bo Wang, Daniele Ramazzotti, Luca De Sano, Junjie Zhu, Emma Pierson, Serafim Batzoglou

## Abstract

**Motivation:** We here present SIMLR (<u>S</u>ingle-cell <u>I</u>nterpretation via <u>M</u>ulti-kernel <u>L</u>ea<u>R</u>ning), an open-source tool that implements a novel framework to learn a cell-to-cell similarity measure from single-cell RNA-seq data. SIMLR can be effectively used to perform tasks such as dimension reduction, clustering, and visualization of heterogeneous populations of cells. SIMLR was benchmarked against state-of-the-art methods for these three tasks on several public datasets, showing it to be scalable and capable of greatly improving clustering performance, as well as providing valuable insights by making the data more interpretable via better a visualization.

**Availability and Implementation:** SIMLR is available on GitHub in both R and MATLAB implementations. Furthermore, it is also available as an R package on bioconductor.org.

**Supplementary Information:** Supplementary data are available at *Bioinformatics* online.

## 1 Introduction

The recent development of high resolution single-cell RNA-seq (scRNA-seq) technologies increases the availability of high throughput gene expression measurements of individual cells. This allows us to dissect previously unknown heterogeneity and functional diversity among cell populations [1]. In this line of work recent efforts (see [2, 3, 4]) have demonstrated that *de novo* cell type discovery of functionally distinct cell sub-populations is possible via unbiased analysis of all transcriptomic information provided by scRNA-seq data. However, such analysis heavily relies on the accurate assessment of pairwise cell-to-cell similarities, which poses unique challenges such as outlier cell populations, transcript amplification noise, and dropout events (*i.e.*, zero expression measurements due to sampling or stochastic transcriptional activities) [5].

Recently, new single-cell platforms such as DropSeq [6] and GemCode single-cell technology [7] have enabled a dramatic increase in throughput to hundreds of thousands of cells. While such technological advances may add additional power for *de novo* discovery of cell populations, they also increase computational burdens for traditional unsupervised learning methods.

To address all of the aforementioned challenges, SIMLR was proposed in [8] as a novel framework that learns an appropriate cell-to-cell similarity metric from the input single-cell data. The learned similarities enable effective dimension reduction, clustering, and visualization of cells. SIMLR provides a more scalable analytical framework, which works on hundreds of thousands of cells without any loss of accuracy in dissecting cell heterogeneity.

## 2 The SIMLR framework

SIMLR is is available in both R and MATLAB implementations. The framework is capable of learning cell-to-cell similarities among gene expression data of individual cells, which have been shown to capture different representations of the data. The approach combines multiple Gaussian kernels in an optimization framework, which can be efficiently solved by a simple iterative procedure. Moreover, SIMLR addresses the challenge of high levels of noise and dropout events by employing a rank constraint and graph diffusion in the learned cell-to-cell similarity [9]. See Figure 1 for an overview of the framework.

**Figure 1:**
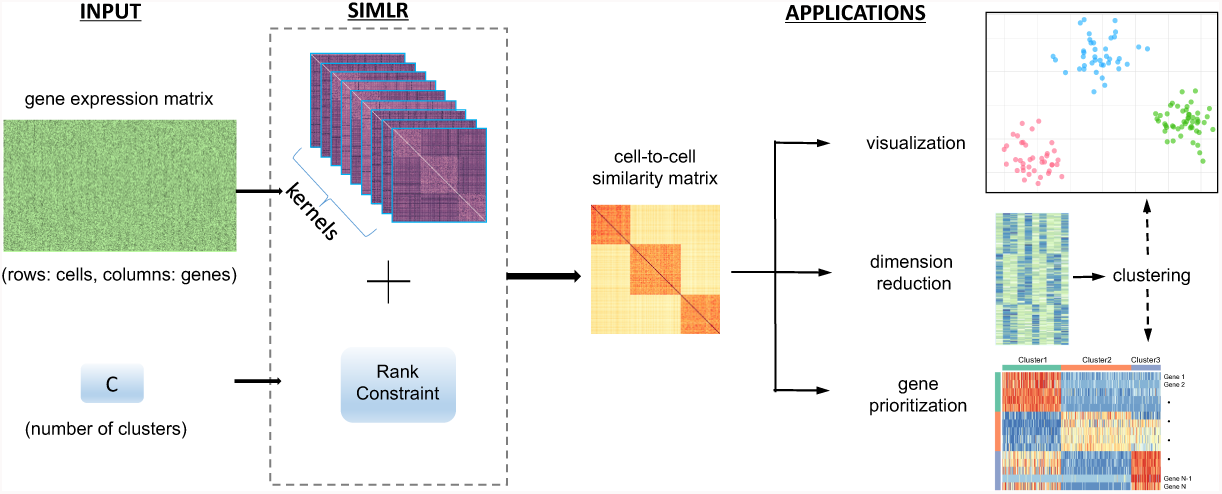
SIMLR pipeline [9]. We start with an input matrix with gene expression observations for a set of genes. SIMLR is then capable of learning a set of cell-to-cell similarities by estimating multiple kernels, with the assumptions of the presence of *C* separable populations within the data. To this extent, SIMLR constraints the similarity matrix to have an approximate block-diagonal structure with *C* blocks where the cells of the same populations to be more similar. The learned similarities can be used for multiple tasks; they can be used to visualize the cells, reduce the dimension of the data, cluster the cells into subgroups and prioritize the most variable genes that explain the differences across the populations.

The framework provides both a standard implementation and a large-scale extension of SIMLR together with two examples to test the methods on the datasets by [10] for the standard SIMLR and [11] for the large-scale extension (see Supplementary Material for details). SIMLR can accurately analyze both datasets within minutes on a single core laptop.

One of the advantages of SIMLR is that the learned similarities can be efficiently adapted into multiple downstream applications. Some applications include prioritizing genes by ranking their concordance with the similarity and creating low-dimensional representations of cells by transforming the input into a stochastic neighbor embedding framework. We refer to the Supplementary Material for detailed use cases of the tool and to [8] for a detailed description of the method and for several applications on genomic data from public datasets.

## 3 Discussion

SIMLR infers the cell-to-cell similarities that are used to perform dimension reduction, clustering, and visualization. While the multiple-kernel learning framework has obvious advantages on heterogeneous single-cell datasets, where several clusters coexist, we also believe that this approach, together with its visualization framework, may also be valuable for data that does not contain clear clusters, such as cell populations that contain cells spanning a continuum or a developmental pathway.

